# *Piromyces struthionis,* sp. nov., a new anaerobic gut fungus from the feces of ostriches

**DOI:** 10.1101/2025.04.22.650011

**Authors:** Kathryn Nash, Julia Vinzelj, Carrie J. Pratt, Mostafa S. Elshahed, Noha H. Youssef

**Affiliations:** Department of Microbiology and Molecular Genetics, Oklahoma State University, Stillwater, OK, USA

**Author notes:** Corresponding Author, Address: 1110 S. Innovation Way, Stillwater, OK, USA. Both authors contributed equally to this study (Kathryn Nash, Julia Vinzelj).

**Keywords:** Fungi, herbivory, *Neocallimastigomycota*, *Aves*

## Abstract

Anaerobic gut fungi (AGF, Neocallimastigomycota) are a clade of basal, zoospore-producing fungi within the subkingdom Chytridiomyceta and known inhabitants of the alimentary tract of animal hosts. To date, 22 genera and 38 species have been described, most originating from herbivorous mammals. Here, we report on the isolation and characterization of a novel species of Neocallimastigomycota from an avian host. Multiple AGF strains were isolated from ostrich feces obtained from a local farm in Oklahoma (USA). All strains formed small, irregular-shaped white colonies with darker centers, displayed a filamentous rhizoidal structure with monocentric thallus developmental patterns, and produced mostly monoflagellated zoospores. The type strain produced terminal sporangia that were predominantly globose, often exhibiting cup-shaped, and occasionally elongated sporangiophores. Sporangiophores characteristically exhibited constrictions at irregular intervals, giving them a pearls-on-a-chain-like appearance. Phylogenetic analysis using the D1-D2 region of the LSU rRNA gene (D1-D2 LSU), ribosomal internal transcribed spacer-1 (ITS1), and RNA polymerase II large subunit (RPB-II) grouped all isolates as a separate species within the genus Piromyces. Transcriptomic analysis indicated an average amino acid identity (AAI) of 80.34 % (± 3.27 %) between the type species and members of the genus Piromyces, and 62.93-76.05 % between the type species and all other AGF taxa outside Piromyces. Based on the morphology, phylogenetic analysis, and AAI values, we propose accommodating these strains as a novel species of Piromyces for which the name Piromyces struthionis is proposed. The type strain for this species is Ost1.

## Introduction

Anaerobic gut fungi (AGF) represent a unique phylum (*Neocallimastigomycota*) within the fungal kingdom. These strictly anaerobic fungi are intrinsic members of the gut microbiome in herbivores, where they play a central role in the degradation of complex carbohydrates in plant biomass (Gruninger et al., 2014). To date, AGF have largely been isolated from mammalian herbivores, especially domesticated ones, e.g. cattle, sheep, goat, horses (Hanafy et al., 2022), with recent increasing reports from wild mammalian hosts (Hanafy et al., 2018, 2020, 2021; Stabel et al., 2020). Twenty of the 22 currently described AGF genera were obtained from mammalian hosts, with the remaining two recently isolated from tortoises (Pratt et al., 2023). Twelve AGF genera were assigned to four families, while ten remain unassigned at the family level (Hanafy et al., 2023). Only five genera (*Anaeromyces*, *Neocallimastix*, *Piromyces*, *Capellomyces*, and *Orpinomyces*) harbor more than one species (Elshahed et al., 2022; Hanafy et al., 2022).

The recent isolation of AGF from tortoises (Pratt et al., 2023) suggests a broader scope for potential AGF hosts. It provides a strong indication of the presence of AGF in other non-mammalian herbivores. Ostriches (*Struthio camelus*) are large, flightless, herbivorous birds in the order *Struthioniformes*. Ostriches are unique amongst *Aves* in possessing a digestive system with an expanded caecum and colon, hindgut fermentation capabilities, and a long feed retention time (22 - 40 hours) (Fritz et al., 2012). Such characteristics render them potential hosts for harboring AGF. Our recent efforts have identified the occurrence of AGF in fecal samples of ostriches (Vinzelj et al., 2025; manuscript currently under review). Here, we report on the isolation and characterization of five AGF strains from the fecal samples of two female ostriches from a farm in Oklahoma (USA). Morphological and phylogenetic analysis indicated an affiliation to the genus *Piromyces*, and we propose a novel species (*P. struthionis,* sp. nov.) for their accommodation.

## Materials and Methods

### Samples

Fresh fecal samples were collected from two female ostriches at Snider Family Exotics farm (Fletcher, OK, USA, 34.757288, -98.152112) in April 2022. The ostriches were fed a diet of corn pellet mix. Samples were collected in 50 mL conical centrifuge tubes as quickly as possible after deposition to ensure minimal exposure to oxygen. They were transferred on ice to the laboratory within 24 h of collection and then frozen at −20 °C. Enrichment and isolation started in June 2024, 26 months after initial collection.

### Isolation

One gram of fecal material was serially diluted into rumen fluid cellobiose (RFC) media (Calkins et al., 2016) supplemented with antibiotics (50 µg/mL each of chloramphenicol, penicillin, kanamycin, and norfloxacin, and 20 µg/mL streptomycin) and 0.1 g/mL switchgrass.

Similarly, enrichments were also started in rumen fluid-free (RFF) medium (Vinzelj et al., 2023) supplemented with antibiotics (20 µg/mL penicillin G sodium salt, 20 µg/mL streptomycin sulfate, and 10 µg/mL ampicillin sodium salt) and 0.1 g/mL switchgrass. All enrichments were incubated at 39°C and showed growth after 19 to 21 days. Dilutions exhibiting signs of fungal growth (e.g., plant material floating and clumping, visible fungal biomass, and production of gas bubbles) were analyzed via light microscopy and subcultured into fresh RFC or RFF media. Roll tubes were prepared from successful enrichment tubes and incubated at 39°C until single colonies were obtained. Three rounds of colony picking and roll tubing were repeated to ensure culture purity. In total, three rounds of enrichments were conducted, resulting in the isolation of five strains, which are maintained at 39 °C by bi-weekly subcultivation in RFC medium.

### Temperature preferences

To determine the minimal, maximal, and optimal growth temperatures for the type strain, the isolate was grown at 30, 32, 35, 39, 41, and 43 °C (five replicates per temperature) in RFC medium in Balch tubes (CLS-4209, ChemGlass Life Sciences, New Jersey, U.S.A.). Growth was determined by both visual inspection (biomass production, sticking of biomass to glass) and measurement of gas pressure accumulation in the headspace using a digital pressure gauge (MediaGauge, SSI Technologies). Media-to-headspace ratio was identical in all experiments (10 mL liquid and 17 mL headspace). A mammalian isolate (*Piromyces* sp. isolate NO 2.2, obtained from a camel fecal sample) was utilized to compare the growth of mammalian and avian *Piromyces* isolates. Negative (uninoculated) controls were also included. Both strains and the negative controls were subcultured five consecutive times for each temperature at four-day intervals.

### Morphological and microscopic characterization

Colony morphology and growth patterns on agar and in liquid RFC media were examined at the optimal growth temperature (39 °C). Microscopic analysis of cultured stained with Lactophenol-Cotton Blue Stain was used to observe zoospores during log phase (1 day after subculturing), while sporangia and hyphae development were observed during the late-log through early stationary phases (2-5 days after subculturing). Both light (Olympus BX51 instrument equipped with an MU503-GS AmScope digital camera), and confocal (Zeiss LSM 980 Airyscan 2 confocal laser scanning microscope) microscopy were used. To examine nuclei localization, samples were stained with (VWR, Cat.No. VW3427-0, discontinued) and 4,6′-diamidino-2-phenylindole (DAPI; 10 µg/mL). Sizes of various microscopic structures were measured using Fiji software (Schindelin et al., 2012).

### Phylogenetic analysis

Fungal biomass was harvested by centrifugation, and DNA was extracted using a DNeasy PowerPlant Pro Kit (Qiagen Corp., Germantown, MD, USA) according to the manufacturer’s instructions. The D1 and D2 regions of the 28S large ribosomal subunit gene (D1-D2 LSU) were amplified using the primer pair NL1 (5′-GCATATCAATAAGCGGAGGAAAAG-3′) and NL4 (5′-GGTCCGTGTTTCAAGACGG-3′). Amplicons were cleaned using a PureLink PCR Purification Kit (ThermoFisher Scientific, Waltham, MA, USA) and subjected to Sanger sequencing at the Oklahoma State University DNA Protein Core Facility (Stillwater, OK, USA). For the type strain, both the ITS1 region and the D1-D2 domains of LSU rRNA gene were amplified, and amplicons were cloned into a pCR-XL-2-TOPO cloning vector according to the manufacturer’s instructions (ThermoFisher Scientific, Waltham, MA, USA) as described previously (Pratt et al., 2023). As recently recommended (Elshahed et al., 2022), twelve clones were Sanger sequenced at the Oklahoma State University DNA Protein Core Facility (Stillwater, OK, USA) or Eurofins Genomics (Louisville, KY, USA) to examine intra-strain variability. ITS1 and D1-D2 LSU from the obtained clones were aligned to reference ITS1 and D1-D2 LSU sequences using MAFFT (Katoh et al., 2019), and the alignment was used to construct maximum likelihood phylogenetic trees in FastTree (Price et al., 2010) using *Chytriomyces* sp. WB235A isolate AFTOL-ID 1536 as an outgroup. Bootstrap values were calculated based on 100 replicates.

For RNA polymerase II large subunit (RPB1) protein trees, amino acid sequences were identified within the isolates’ transcriptome’s predicted peptides (see below) and aligned using MAFFT (Katoh et al., 2019). IQ-TREE (Minh et al., 2020) was used to predict the best substitution model and to generate maximum-likelihood trees under the predicted best model. Options ‘–alrt 1000’, ‘-abayes’, and ‘–bb 1000’ were added to the command line to perform the Shimodaira–Hasegawa approximate-likelihood ratio test (SH-aLRT), approximate Bayes tests, and ultrafast bootstrap (UFB), respectively. IQ-TREE analysis resulted in the generation of phylogenetic trees with three support values (SH-aLRT, aBayes, and UFB) on each branch.

### Transcriptomic sequencing and AAI determination

Cultures were grown in RFC media to late exponential phase, early log-phase (4-5 days) and vacuum filtered. Total RNA was extracted using the Macherey-Nagel™ NucleoSpin™ RNA Mini kit according to the manufacturer’s instructions. Total RNA was used for RNA-seq on an Illumina NextSeq 2000 platform using a 2×150 bp paired-end library at the One Health Innovation Foundation lab at OSU. Trinity (version 2.6.6; Grabherr et al., 2011; Haas et al., 2013) was used for quality trimming and *de novo* assembly of RNA-seq data using default parameters. Redundant transcripts were clustered with an identity parameter of 95 % (–c 0.95) using CD-HIT (Fu et al., 2012). Peptide and coding sequence predictions were conducted using TransDecoder (version 5.0.2; https://github.com/TransDecoder/TransDecoder) with a minimum peptide length of 100 amino acids. The predicted peptides were used for average amino acid identity (AAI) calculations, as well as for extracting the single-copy protein RPB1 for phylogenetic assignment (see above). For AAI calculations, we included predicted peptides from other AGF transcriptomes (*n* = 60; Pratt et al., 2024). AAI values were calculated for all possible pairs in the dataset using the *aai.rb* script available as part of the Enveomics collection (Rodriguez-R and Konstantinidis, 2016).

### Comparative gene content analysis

Predicted peptides from ten *Piromyces* transcriptomes (strains isolated from mammalian hosts; (Pratt et al., 2024) were compared to the predicted peptides from *Piromyces struthionis* (Strain Ost1). Peptides were classified against COG, KOG, and KEGG classification schemes as previously described (Pratt et al., 2024). In addition, the overall CAZyme content was predicted using run_dbcan4 (https://github.com/linnabrown/run_dbcan).

### Sequence and data deposition

RNA-seq reads were deposited in NCBI SRA under BioProject accession number PRJNA1231060. Clone sequences of the D1-D2 region of the LSU rRNA were deposited in GenBank under accession numbers PV213533-PV213569. Clone sequences of the ITS1 region were deposited in GenBank under accession numbers PV213570-PV213582.

## Results

### Isolation

Five strains (Ost1, OK2.2, OJ2.2, OJ2.4, OSF2.3.3.1) were isolated from fecal samples obtained from two different ostriches in five separate enrichment procedures (Table 1). All strains displayed identical colony morphology, microscopic features, and D1-D2 LSU sequences (sequence divergence 99.17±0.02%). Strain Ost1 was selected for detailed characterization and designated as the type strain.

**Table 1:** Summary of the successful isolations from ostrich fecal samples.

| Strain | Fecal sample | Media* | Enrichment bottles | Incubation temperature (°C) |
| --- | --- | --- | --- | --- |
| Ost1 | OSF1 | RFF+SG | Balch tubes with 10 mL media and 1 g of feces | 39 |
| OJ2.2 | OSF1 | RFF+SG | Balch tubes with 10 mL media and 1 g of feces | 39 |
| OJ2.4 | OSF1 | RFF+SG | Bottles with 50 mL media and 5 g of feces | 39 |
| OK2.2 | OSF1 | RFF+SG | Balch tubes with 10 mL media and 1 g of feces | 41 |
| OSF2.3.3 | OSF2 | RFC + MCC | Balch tubes with 10 mL media and 1 g of feces | 39 |

\* RFC = rumen fluid cellobiose medium(Calkins et al., 2016), RFF = rumen fluid free medium (Vinzelj et al., 2023) containing xylan; SG= Switchgrass (0.1 g mL<sup>-1</sup>), MCC = microcrystalline cellulose (0.1 g mL<sup>-1</sup>)

### Macroscopic and microscopic growth characteristics

On agar roll tubes, strain Ost1 exhibits small (0.5 - 1 mm) colonies that are light in color, with solid centers and lighter edges (Figure 1A). In liquid RFC medium, biomass creates thin flakes or a biofilm that tightly adheres to the walls and bottom of the tube (Figure 1B-C). Strain Ost1 produced globose zoospore (Fig. 1D-G), with an average diameter (± SD) of 5.9 ± 1.1 μm (n = 32, range: 3.5-7.8 μm). The majority (approximately 78%) of zoospores were monoflagellated (Figure 1D) with an average flagellum length (± SD) of 16.3 ± 5.9 μm (n = 40, range: 10.2-21.7 μm). Approximately 13% of zoospores were biflagellated (Figure 1E), and ∼9% were triflagellated (Figure 1F). Tetra-flagellated zoospores were exceedingly rare (Figure 1G). Occasionally, multiple zoospores were observed clumping together (Figure 1H-I)

**Figure 1.**
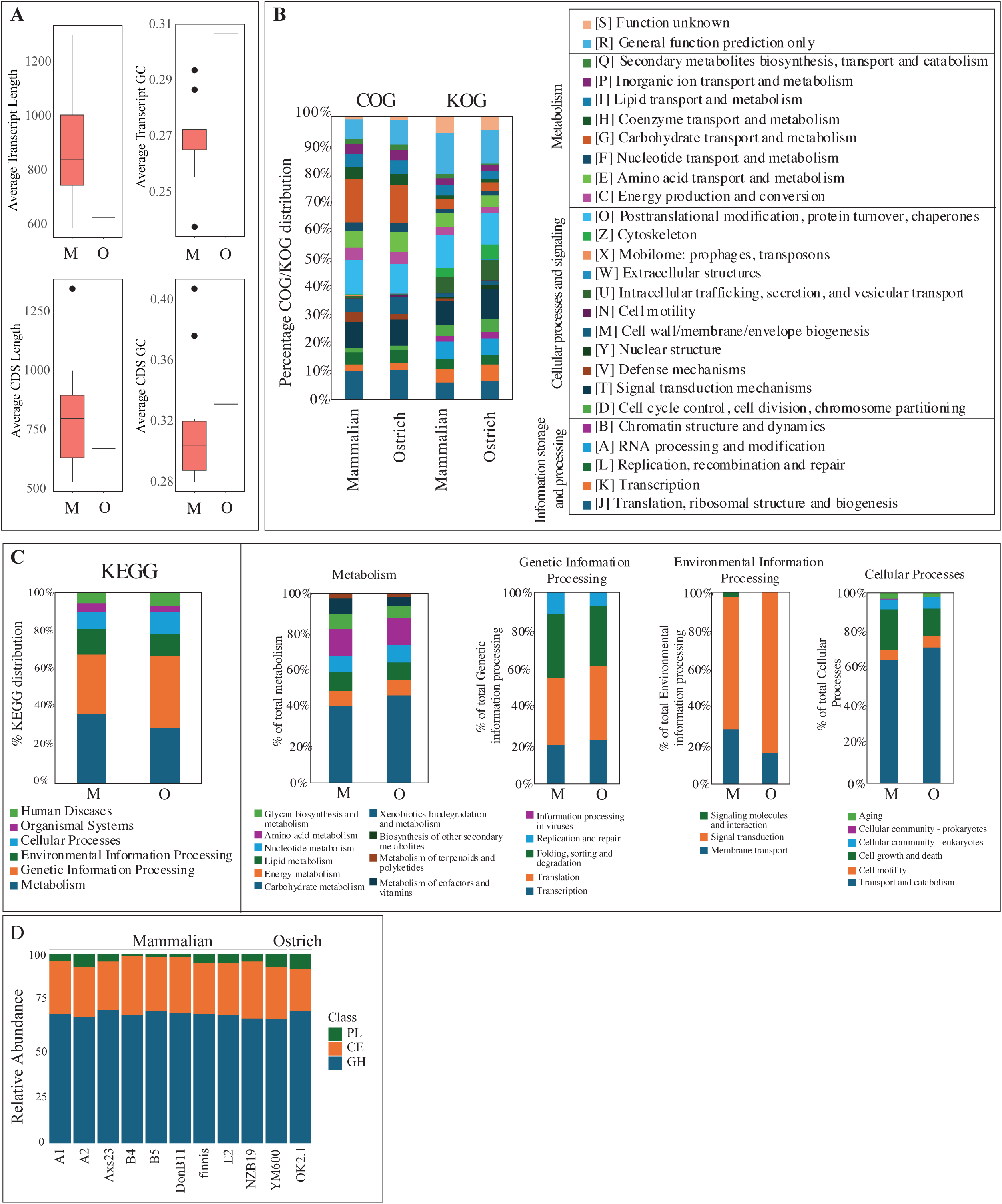
Macroscopic and microscopic characteristics of *Piromyces struthionis* strain Ost1 observed via light (D-V, X-Z) and confocal (W & AA) microscopy. On roll tubes (A), *P. struthionis* exhibits small (0.5 - 1 mm) round colonies that are light in color with solid centers due to extensive filamentous growth and lighter edges where less growth has occurred. In liquid RFC medium (B-C) biomass lightly sticks to itself and firmly attaches to the walls of the tube. Zoospores (D – I) are monoflagellated with one (D), two (E), three (F), or four (G) flagella, and occasionally cluster together (H-I). Encysted zoospores with no flagella (J), leading to double (arrows) (K), and single (L-M) germ tube production. Note the branching of the germ tube in (M). Monocentric thallus with nuclei only in the sporangia (N-O). Both endogenous (P) and exogenous (Q-V) sporangia developed, with exogenous development observed more frequently. Exogenous sporangia developed at the end of a sporangiophore that often exhibited a cup shape (Q-S), although wide flattened sporangiophores were also observed (T). Many sporangiophores showed multiple constrictions giving the appearance of beads-on-a-string (S-U) were very characteristic of *Piromyces struthionis*. Very long sporangiophores were also observed (V) with occasional subsporangial swelling forming at the end of the sporangiophore (arrow in V). Sporangia are typically globose (N-W) but can also be elongated (arrows in W). The rhizoidal system (X, Y) was often hard to observe due to the thickness of the sporangial mats. Rhizoids were thin with minimal branching. Zoospore release was rarely observed and occurred either through an apical pore (Z), or via rupturing of the sporangia (AA). However, these could be artefacts during slide preparation/handling.

Zoospore encystment (Figure 1J) was occasionally observed. Germ tubes and branching rhizoidal systems (Figure 1L-M) were observed arising from one end of the encysted zoospore. Encysted zoospores with two germ tubes were also observed giving rise to pseudo-intercalary like sporangia (Figure 1K).

A monocentric thallus developmental pattern was observed, with nuclei not migrating to the rhizoids (Figure 1N, O). Both endogenous (where the zoospore cyst enlarges and develops into the sporangium, Figure 1P), and exogenous (where the zoospore cyst germinates bipolary, with rhizoids developing at one end of the zoospore cyst, and a sporangiophore developing at the opposite end with the sporangium forming at the end of the sporangiophore, Figure 1Q-V) sporangial developmental patterns were observed. The majority of sporangia developed exogenously (Figures 1N, Q-V). Sporangia were mostly globose (Figure 1W), though a few elongated sporangia were identified (arrow in 1W). A diameter of 28.4 ± 11.9 μm (average ± SD) was observed (n = 51).

Sporangiophores in strain Ost1 were mostly eggcup shaped (Figure 1Q-S), but wide-flattened sporangiophores (Figure 1T), and extremely elongated sporangiophores were also encountered (Figure 1R, S, V). The average length of all recorded sporangiophores (n = 12) was 39.1 ± 12.7 μm (average ± SD). Sub-sporangial swellings (apophysis) were occasionally observed (arrow in 1V). Long and multi-constricted sporangiophores were frequently observed (Figures 1S, U), sometimes ending in an egg-cup shape (Figure 1S). On the edges of the sporangial mats, mostly thin rhizoids were observed with minimal branching (Figure 1X, Y).

Zoospore release was not frequently identified. However, two mechanisms were observed: an apical pore (Figure 1Z) and rupturing of the sporangial wall (Figure 1AA). However, both observations could have been an artefact during slide preparation/handling.

### Temperature preferences

Strain Ost1 grew at a relatively wide range of temperatures (32-41 °C) in the initial subculture, and the growth was maintained for five subsequent subcultures (n = 5). The strongest growth was between 35-39 °C, and optimal growth occurred at 39 °C. In comparison to mammalian *Piromyces* strain NO2.2 (obtained from a camel), *Piromyces* strain Ost1 was more tolerant to lower temperatures (32 °C and 35 °C). Both, however, were unable to sustain growth at 30 °C. At higher temperatures, no significant difference in tolerance between the two strains was observed, and both grew very weakly at 43 °C (Figure 2).

**Figure 2.**
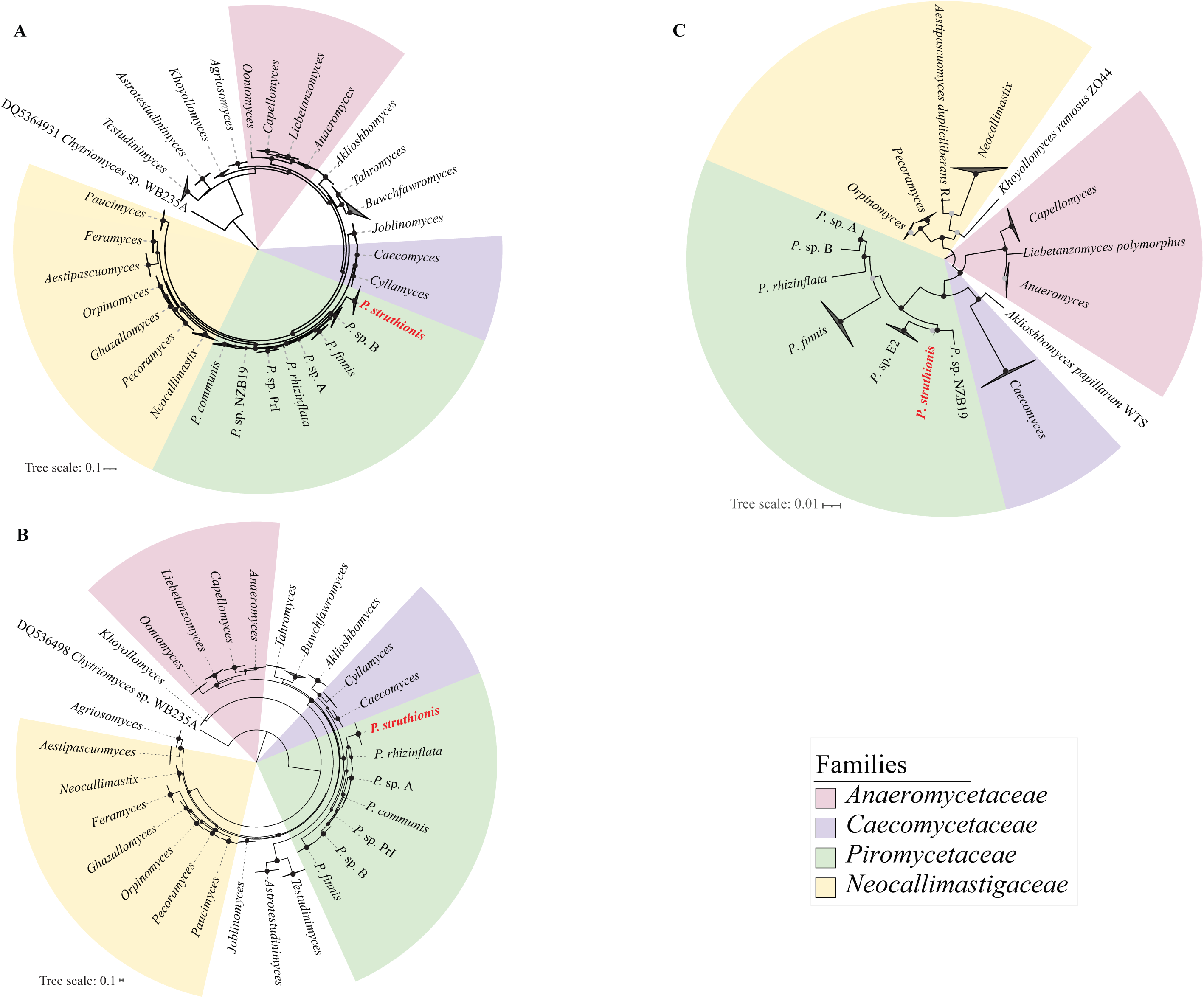
Temperature growth preferences of *Piromyces struthionis* strain Ost1 and *Piromyces* sp. NO2.2 (isolated from camel feces). Gas pressure in PSI (as a proxy for growth) is shown on the y-axis. Strains were grown in five replicates at different temperatures (X-axis) and subcultured every 4 days. The gas pressure buildup was measured at the end of each subculture. Averages ± standard deviation of all 5 subcultures from all five replicates at each growth temperature are shown for PSI values.

### Phylogenetic analysis

Phylogenetic analysis using the D1-D2 region of the LSU rRNA placed strain Ost1 within the *Piromyces* clade with high bootstrap support (Fig. 3A). Sequence divergence within Ost1 clones was extremely low ranging between 0-0.24% (average 0.055±0.044). The closest *Piromyces* isolate to strain Ost1 was to a *Piromyces* sp.A1 (MT085684.1) isolated from cattle. Sequence similarity of *Piromyces* Ost1 clones to clones from *Piromyces* sp.A1 ranged between 95.52-98.17 (average, 97.35±0.52%) sequence similarity.

**Figure 3.**
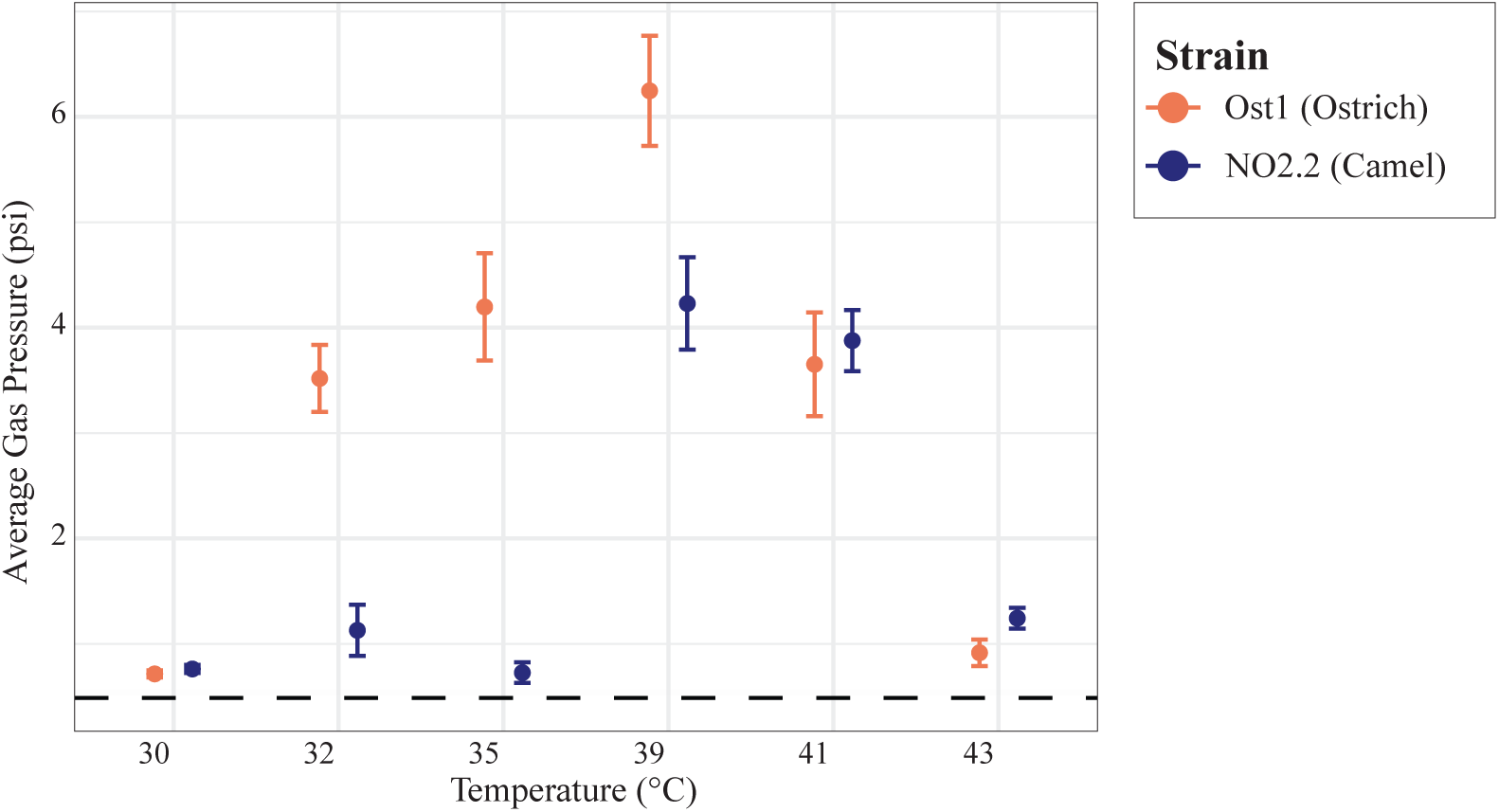
Phylogenetic analysis of *Piromyces struthionis* using D1-D2 LSU (a), ITS1 (b), and RPB1 (c) as phylogenetic markers. Alignments were created in MAFFT (Katoh et al., 2013). Trees in (a) and (b) were reconstructed using the maximum-likelihood approach implemented in FastTree (Price et al., 2010) using *Chytriomyces* sp. WB235A isolate AFTOL-ID 1536 as an outgroup. The RPB1 tree in (c) was constructed in IQ-TREE (Minh and others 2020). Bootstrap values in (a) and (b) were calculated on the basis of 100 replicates and are shown for nodes with >50 % support as black spheres. The three bootstrap support values in (c) (SH-aLRT, aBayes, and UFB) are shown as colored dots as follows: all three support values >50 %, black; 2/3 support values >50 %, dark grey; 1/3 support values >50 %, light grey. Scale bars indicate the number of substitutions per site.

Phylogenetic analysis using the ITS1 locus also placed Ost1 within the *Piromyces* clade with high bootstrap support (Fig. 3B). Sequence divergence within Ost1 clones was extremely low (0.049±0.054%). The closest relative was *Piromyces rhizinflata* strain SFH682 (MK775329.1) isolated from sheep (82.53-84.53%, average 83.67±0.51%).

Finally, phylogeny based on RPB1 marker also placed strain Ost1 within the *Piromyces* clade with the closest *Piromyces* relative being *Piromyces* sp. E2 (Fig. 3C).

### AAI analysis

Compared to other *Piromyces* species with sequenced transcriptomes (n=10), Strain Ost1 exhibited pairwise AAI values ranging from 73.63% (to *Piromyces* sp. B strain B5) to 84.24% (to *Piromyces* sp. NZB19 strain Ors32), with an average AAI of 80.33±3.47%. In contrast, lower AAI values were observed between Ost1 and all other AGF genera, i.e. members of the families *Anaeromycetaceae* (73.88±0.87%), *Caecomycetaceae* (74.52±0.75%), and *Neocallimastigaceae* (73.98±0.70%), as well as to various *incertae sedis* genera (e.g., *Khoyollomyces* 71.57%, *Aklioshbomyces* 76.05%, *Astrotestudinimyces* (63.17±0.34%) and *Testudinimyces* (64.41±0.58%)) (Table 2).

**Table 2.**
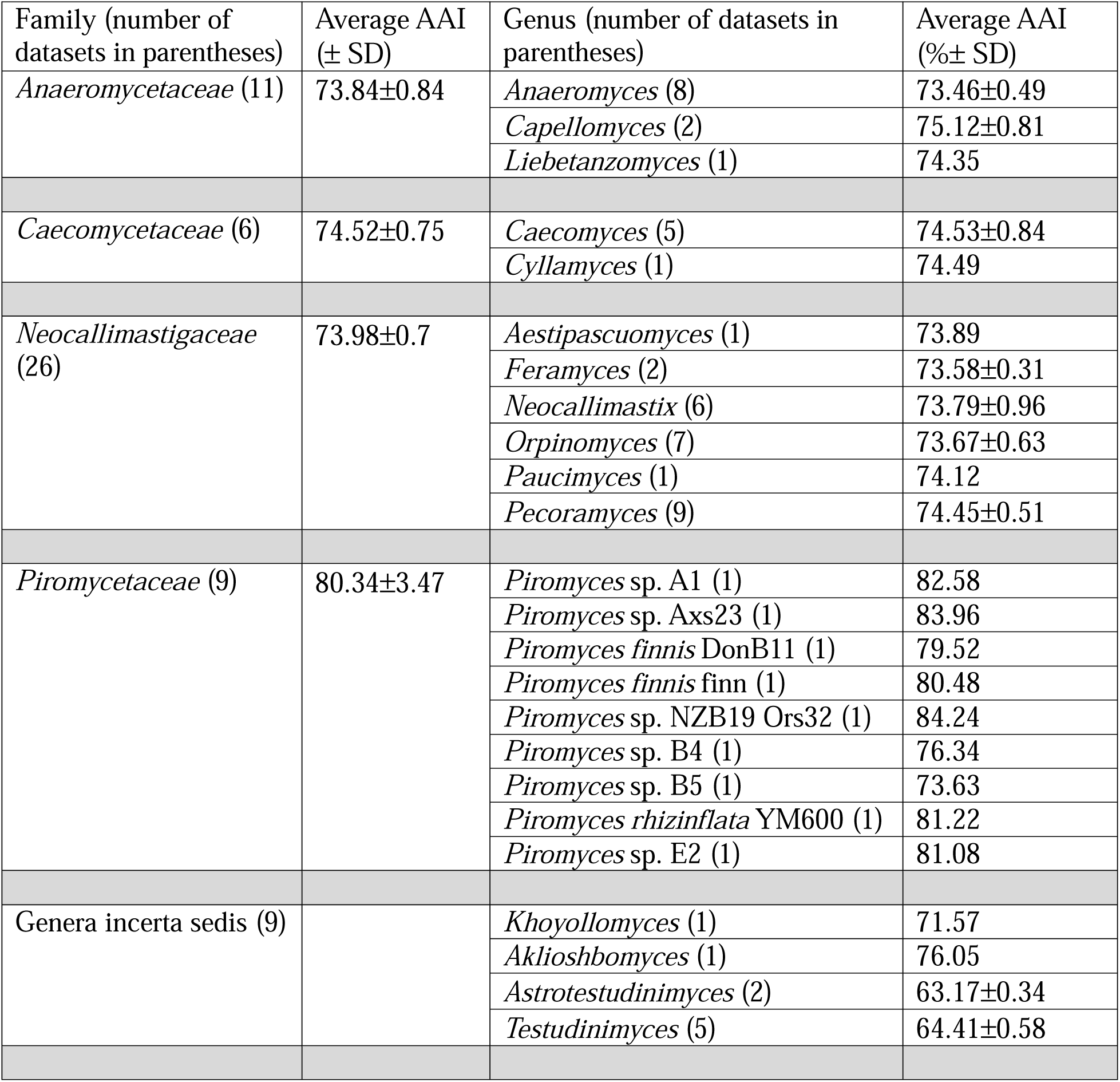
Average amino acid identity values comparing *Piromyces struthionis* OST1 to other AGF genera.

### Comparative gene content analysis

Comparative analysis of the Ost1 transcriptome to the 10 *Piromyces* transcriptomes originating from mammalian isolates showed slightly shorter transcript length and higher GC content in both the transcript and the coding sequence (albeit with no significant difference due to the lack of ostrich isolate replicates) (Figure 4A). The overall COG, KOG, and KEGG composition did not vary significantly by the source of isolation (mammalian versus ostrich) (Figure 4B-C). Similarly, the broad makeup of the CAZyome (relative abundances of total GHs, CEs, and PLs) was very similar in all *Piromyces* strains (Figure 4D, Table S1).

**Figure 4:**
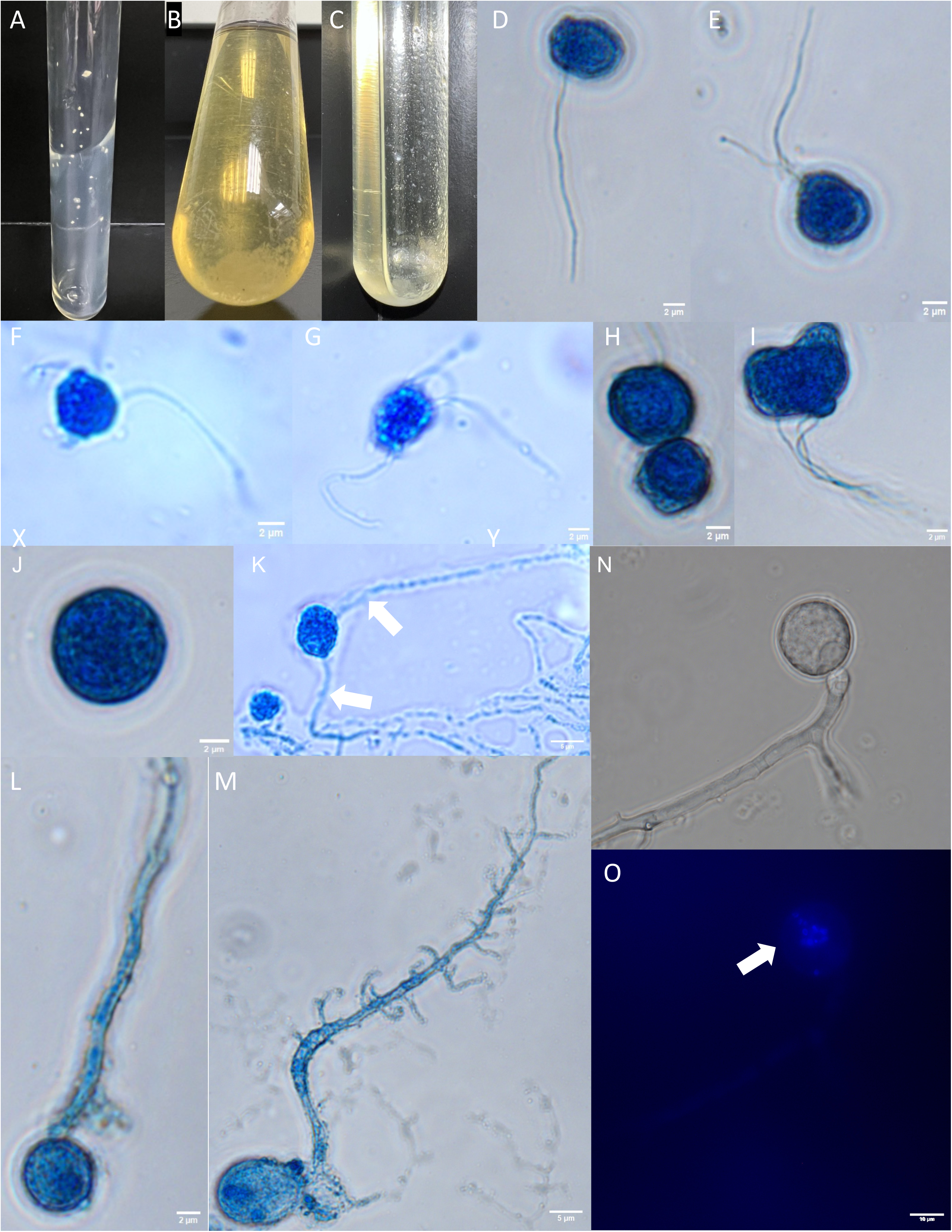
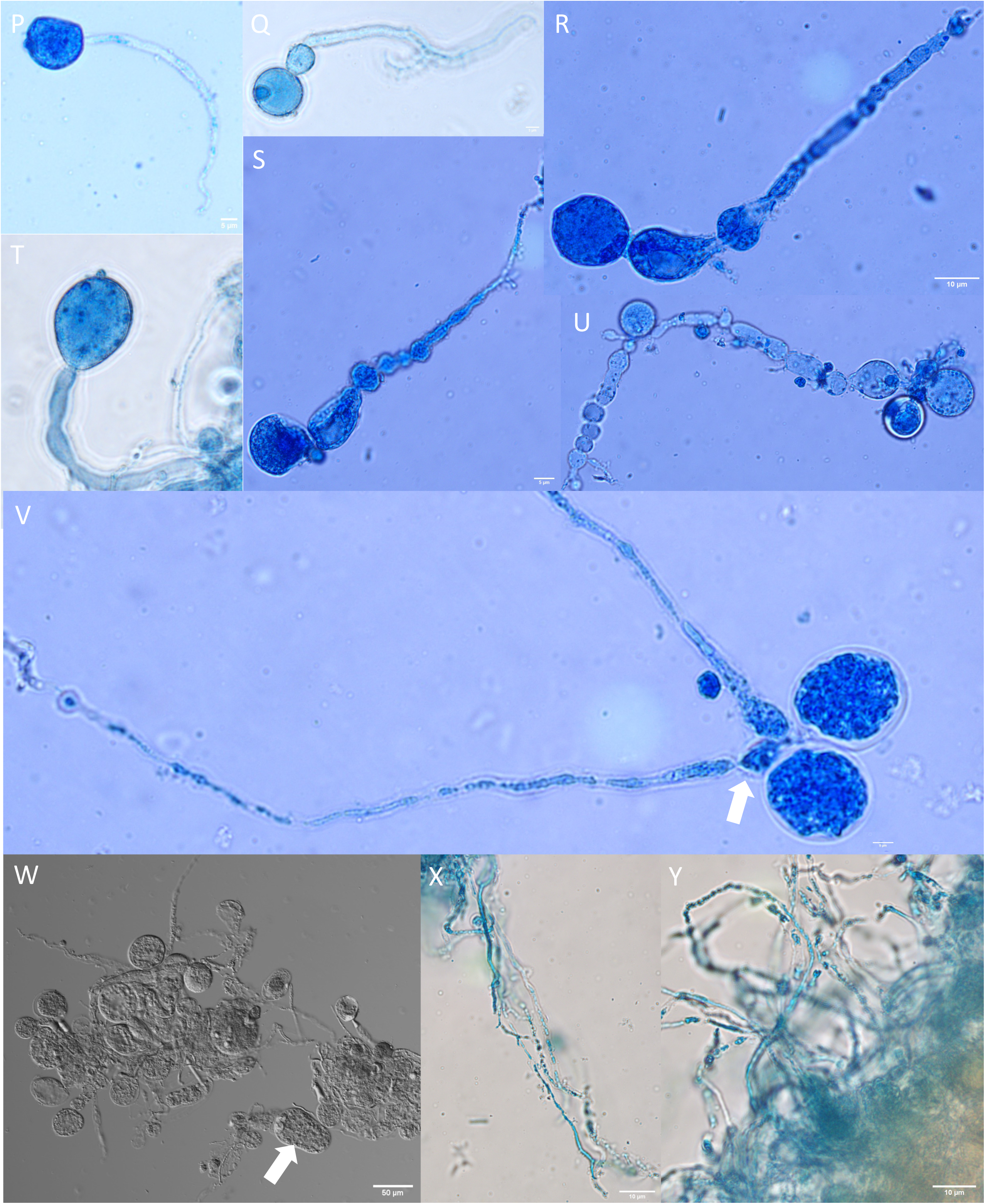
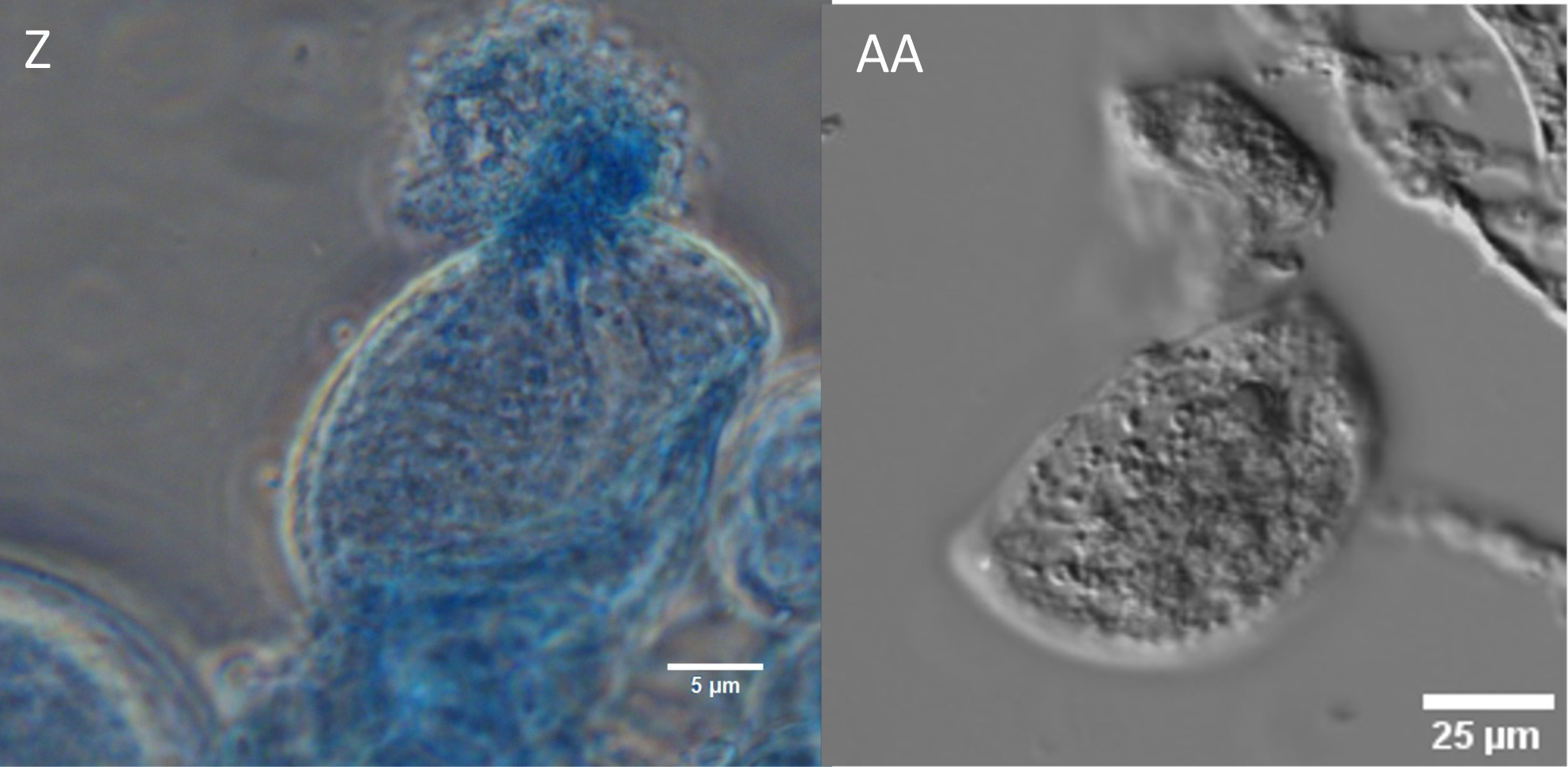
Comparative general features and total gene content analysis of *Piromyces struthionis* strain Ost1 transcriptome (O) to 10 other *Piromyces* transcriptomes obtained from mammalian hosts (M). **(A)** Distribution of transcript length (top left), transcript GC content (top right), coding sequence length (bottom left), and coding sequence GC content (bottom right) for the mammalian *Piromyces* transcriptomes (M) versus the *Piromyces struthionis* strain Ost1 transcriptome (O). Values for mammalian *Piromyces* transcriptomes are shown as a boxplot. ANOVA for pairwise comparison was not possible due to the lack of replicates of ostrich *Piromyces* transcriptomes. **(B-C)** Total gene content comparison between mammalian sourced (left stacked columns, M) and ostrich sourced (right stacked columns, O) transcriptomes using COG/KOG **(B)**, and KEGG **(C)** classification. KEGG classification in (C) is further broken down into the four main categories: Metabolism, Genetic Information Processing, Environmental Information Processing, and Cellular Processes. **(D)** CAZyome comparison of *Piromyces struthionis* strain Ost1 transcriptome to 10 other *Piromyces* transcriptomes obtained from mammalian hosts. Relative abundance of CEs, GHs, and PLs in the compared transcriptomes. Mammalian transcriptomes are shown first, followed by *Piromyces struthionis* strain Ost1. For the detailed distribution of all CAZy families, refer to Table S1.

### Taxonomy

*Piromyces struthionis* Kathryn Nash, Julia Vinzelj, Mostafa S. Elshahed, & Noha H. Youssef., sp. nov.

Mycobank number: MB858494

*Typification*: U.S.A. OKLAHOMA: Stillwater, 36.12′N, 97.06′W, ∼300 m above sea level, isolated from freshly deposited, then frozen feces of an ostrich (*Struthio camelus*). Holotype material from the culture of isolate Ost1 (3-day-old cultures, killed and preserved in 5% glutaraldehyde) is stored at the Oklahoma State University Culture Collection. Ex-type culture Ost1 type culture is stored cryogenically in liquid nitrogen at the Oklahoma State University Culture Collection. The strain is also maintained in active culture through bi-weekly subcultivation on RFC media. GenBank accession no. PV213570-PV213582 (ITS1), and PV213533-PV213569 (28S rRNA, partial sequence; D1-D2 regions).

*Etymology*: The species epithet reflects the fact that this fungus was first isolated from ostrich (*Struthio camelus*) feces.

An obligately anaerobic fungus with determinate monocentric thallus and single terminal sporangia. Anucleate rhizoidal system, with minimal branching and constrictions. Mature sporangia are globose or occasionally elongated, with a diameter of (± SD) 28.4 ± 11.9 μm. Sporangiophores unbranched, with an average length of (± SD) 39.1 ± 12.7μm, often forming an eggcup-like or wide-flat swelling below the sporangium. Very long sporangiophores with constrictions are frequently observed. Zoospores formed abundantly, spherical [(± SD) 5.9 ± 1.1 μm diam] with a single flagellum [(± SD) 16.3 ± 5.9 μm long]. Occasional (∼13%) biflagellate zoospores are formed, less frequently triflagellate (∼9%), and tetraflagellate zoospores. Colonies grown on cellobiose exhibit a smooth biofilm-like growth and form small circular colonies (1 mm diam) on agar roll tubes. The clade is defined by the sequences PV213570-PV213582 (ITS1) and PV213533-PV213569 (28S rRNA, partial sequence; D1-D2 regions).

Additional specimens examined: OK2.2, OJ2.2, OJ2.4, and OSF2.3.3.1 U.S.A. OKLAHOMA: Stillwater, 36.12′N, 97.06′W at ∼300 m above sea level, isolated from freshly deposited and immediately frozen feces of ostriches (*Struthio camelus*), in June 2024 by Kathryn Nash and Julia Vinzelj.

## Discussion

In this study, we describe the first anaerobic gut fungal strain isolated from an avian host (the common ostrich, *Struthio camelus*). Ostriches are large, flightless members of the *Palaeognathae* (an infraclass that also includes rheas, cassowaries, emus, and kiwis) that primarily eat grasses, shrubs, and succulents (Fritz et al., 2012; Williams et al., 1993). Compared to other members of the *Palaeognathae*, ostriches have evolved sacculated ceca (one cecum on each side of the small intestine) and a very long, sacculated colon (El-Wahab et al., 2021). The combination of complex lignocellulosic material in their diet, well-developed designated fermentation chambers in their gut, and the long digesta retention times (30-40 hours) (Frei et al., 2015; Mackie, 2002), render them a suitable host for harboring the zoosporic, lignocellulosic-degrading *Neocallimastigomycota*.

Strain Ost1 exhibited monocentric thallus development, filamentous rhizoids, and mainly produced monoflagellated zoospores. These features are defining characteristics for all members of the genus *Piromyces* as well as ten additional AGF genera. More detailed microscopic analysis revealed elongated sporangiophores that originate with no clear distinction from the rhizomycelia and exhibit constrictions that give them a beads-on-a-string-like appearance (Figure 1U). Those characteristics were observed when growing strain Ost1 on various media compositions (RFC and RFF media) and at different temperatures (32-41 °C), and hence could be regarded as stable morphological characteristics. However, the size and shape of the sporangia, the sporangiophore, and the hyphae have previously been reported to depend on the choice of media (Barr et al., 1989; Gold et al., 1988), the choice of C-source (Barr et al., 1989; Joshi et al., 2018), and the age of the culture (Barr et al., 1989; Ho & Barr, 1995). Given the practical impossibility of assessing a strain morphology under all theoretically feasible growth parameters, care should always be taken when using finer microscopic features for species-level identification. Indeed, comparing the finer morphological details of strain Ost1 to other AGF strains revealed multiple similarities within and outside of the genus *Piromyces*. For example, the beads-on-a-string-like sporangiophores were also observed in *Piromyces minutus* (Ho, 1993a), while elongated sporangiophores were reported in *Piromyces minutus*, *Capellomyces foraminis*, and *Aklioshbomyces papillarum* (Ho, 1993b; Hanafy et al., 2020).

For the genus *Piromyces* multiple species have been described from mammalian hosts (*P. communis*, *P. rhizinflata, P. mae, P. dumbonicus, P. minutus, P. spiralis, P. citronii,* and *P. finnis*), but sequence data is only available for three of those species (*P. communis*, *P. rhizinflata,* and *P. finnis)*. Additionally, sequence data is available for other putative *Piromyces* species that are yet to be typified and characterized (*P.* sp. A, *P.* sp. B, *P.* sp PrI, and *P.* NZB19, Figure 3). Our sequence analysis suggests that strain Ost1 is phylogenetically distinct from all *Piromyces* isolates for which sequence data is available. Because phylogenetic comparison between Ost1 and *Piromyces* species for which no sequence data is available (*P. mae, P. dumbonicus, P. minutus, P. spiralis,* and *P. citronii*) is not possible and morphological analysis is unreliable, we cannot conclusively investigate whether strain Ost1 is a representative of *P. mae, P. dumbonicus, P. minutus, P. spiralis,* or *P. citronii*. However, the ecological distribution of strain Ost1 (Vinzelj et al., 2025) shows it to be almost exclusively present in ostrich feces, and either very rare or absent in mammalian feces. This host specificity is in stark contrast to the ubiquity of various mammalian-sourced *Piromyces* species, which typically colonize a wide range of mammalian hosts and niches such as the mammalian rumen, hindgut and caecum.

Based on the differences in marker sequence regions (ITS, LSU, RPBI) (Figure 3), and host specificity, we argue that strain Ost1 is distinct from all other mammalian *Piromyces* species currently described. Furthermore, it should be noted that the genus *Piromyces* in general occupies a unique position within the *Neocallimastigomycota*. Historically, all strains with filamentous rhizoids, monocentric thallus development, and monoflagellated zoospores were assigned to the genus *Piromyces*. However, subsequent phylogenetic assessment based on multiple loci (ITS-1 and LSU) demonstrated the polyphyletic nature of this combination of phenotypes. Multiple genera that are morphologically similar but phylogenetically unrelated to the original *Piromyces* clade have been described in the last decade (Callaghan et al., 2015; Dagar et al., 2015; Hanafy et al., 2017, 2020). Recent large-scale phylogenetic studies showed that the molecular boundaries (AAI, ITS1 and D1-D2 LSU) circumscribing the genus *Piromyces* are much broader than those in all other AGF genera (Hanafy et al., 2023). However, due to the history of rank assignment in *Neocallimastigomycota,* the genus *Piromyces* was retained, and assignment to the genus *Piromyces* was reserved for monoflagellated, monocentric, filamentous strains that form a monophyletic clade around the type species (*P. communis*) (Hanafy et al., 2022). Consequently, the molecular boundaries (AAI, ITS1 and D1-D2 LSU) circumscribing the genus *Piromyces* now exceed recommended criteria for genus-level rank assignment within the *Neocallimastigomycota* (Elshahed et al., 2022). Indeed, sequence divergence for strain Ost1 could justify proposing a new genus for its accommodation using the criteria described in Elshahed et al. (2022). Nevertheless, given the historic context for the genus and the lack of clear morphological differences that could justify splitting this expansive genus, we propose accommodating strain Ost1 as a new *Piromyces* species until further evidence arises.

In conclusion, our efforts resulted in the first reported AGF isolate from an avian host. Based on extensive comparative analyses we propose to accommodate the isolate as a novel species within the genus *Piromyces*, *P. struthionis*. We further posit that non-mammalian herbivores represent a relatively untapped reservoir of novel AGF diversity, and that similar efforts could lead to the identification and isolation of multiple yet-unrecognized novel taxa.

## Acknowledgements

We thank the Oklahoma City Zoo, Happy Acres Ostrich Ranch LLC, Snider Family Exotics, and Haley Anthony for providing fecal samples. We would further like to thank Kale Goodwin for his efforts in isolating AGF from ostrich fecal samples and the Boren Veterinary Medical Teaching Hospital for providing rumen fluid.

## Data availability statement

The sequence data supporting the findings of this study are openly available in NCBI SRA (BioProject accession number PRJNA1231060) and GenBank (accession numbers PV213533-PV213582). The authors further confirm that the data supporting other findings of this study are available within the article and its supplementary materials.

