## Supplemental Table 1 for "*Piromyces struthionis,* sp. nov., a new anaerobic gut fungus from the feces of ostriches"

* Equal contribution

**Supplementary Tables**

**Table S1.** Number of transcripts assigned to each of the CAZy families in the eleven *Piromyces* transcriptomes compared in this study.

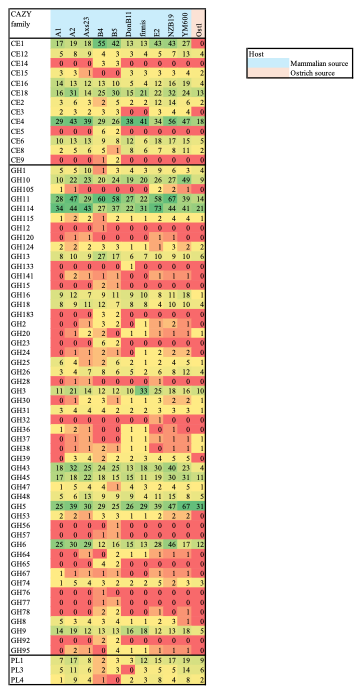
